# Mechanisms of sexually dimorphic adipose tissue remodelling upon prolonged breastfeeding

**DOI:** 10.1101/2025.09.05.674416

**Authors:** Damien Dufour, Sandra Kristine Stølen Bryne, Florian Dapsance, Noemi Ferrito, Ingrid Plotton, Knut Tomas Dalen, Philippe Collas, Charlotte Boccara, Nolwenn Briand

## Abstract

Early developmental cues exert lasting influence on the adipose tissue function and the metabolic health throughout adulthood. In mice, adipose tissue remodelling during the weaning transition programs long-term tissue plasticity, setting its capacity for expansion and thermogenesis in response to environmental stimuli. However, the mechanisms driving this remodelling, and the role of weaning-associated dietary changes in this process, are unknown. Here, we characterise the emergence of sexually dimorphic patterns of adipose tissue distribution and function during the post-weaning period. As subcutaneous adipocytes acquire a white phenotype, the tissue microenvironment undergoes a coordinated remodelling, including reduced innervation and an increase in immune cell content. Using a model of prolonged breastfeeding, we show that it promotes adipose tissue expansion through sex-dependent mechanisms, stimulating progenitor proliferation in males and adipocyte hypertrophy in females. In males, this is accompanied by accelerated tissue whitening, which we link to enhanced clearance of extracellular noradrenaline by mature adipocytes. Our findings reveal sex- and diet-dependent mechanisms governing adipose tissue maturation and highlight how early life nutritional cues shape the microenvironment and function of mature adipocytes.

**Highlights:**

- The peri-weaning period associates with sexually dimorphic remodelling of the adipose tissue
- Whitening of the adipose tissue associates with immune infiltration and decrease innervation in both sexes
- Prolonged breastfeeding prevents drop in proliferation in males
- Whitening is accelerated in males during prolonged breastfeeding

## Introduction

The first years of life are essential for long-term metabolic health. Among various line of evidence, studies have shown that extended breastfeeding supports healthy body mass index (BMI)^1^, and that low sugar intake during early childhood is associated with a reduced risk of developing metabolic diseases^2^. During this critical period, the adipose tissue emerges as a key organ whose development and function are pivotal for shaping long-term metabolic trajectories. The early-life phase is particularly dynamic, with the adipose tissue undergoing significant modifications of its cellularity, metabolism and functions^3,4^. Yet, we still know surprisingly little about the physiological mechanisms that drive this postnatal adipose tissue remodelling and the extent of sex dimorphism in these processes. Gaining this knowledge would be crucial to understand pathological adipose expansion and to identify therapeutic targets for both the prevention and the treatment of metabolic disorders such as obesity and diabetes.

The adipose tissue is a highly plastic organ that adapts to and controls energy status. Excess energy is accommodated by adipose tissue expansion, either though the differentiation of new adipocytes via adipogenesis or through the enlargement of existing adipocytes. On the other hand, when energy is needed, white adipocytes mobilise stored lipids through lipolysis, while brown and beige adipocytes dissipate energy as heat via thermogenesis. Although these functions are broadly conserved across mammalian species, their relative contribution to energy homeostasis varies with age and can be altered in pathological conditions such as obesity. For instance, beige adipocytes arise within human subcutaneous adipose tissue during the last intrauterine semester, then gradually disappear with age^5^. A similar remodelling of subcutaneous adipose tissue occurs in mice during the first five weeks after birth^6,7^, underscoring the value of the mouse model for studying early-life adipose biology.

In mice, inguinal subcutaneous WAT (iWAT) is mainly composed of white adipocytes until postnatal day 10 (p10), when beige adipocytes emerge in a leptin dependant manner^8^. By postnatal day 21 (p21), beige adipocytes represent the predominant subtype^6,7,9,10^. At this stage, the tissue transitions back to a white phenotype (whitening) and undergoes a surge in adipose progenitor proliferation^11^. A similar remodelling occurs in other depots such as retroperitoneal^9^ and perirenal WAT^12^. Conversely, the epididymal WAT remains composed of white adipocytes throughout this period^13^. The mechanisms triggering such remodelling remain unclear. It was established that this period is particularly sensitive to environmental changes, as a 2-week-long high fat diet between p21 and p35, leads to long-term insulin desensitisation^11^. Moreover, these transiently beige adipocytes form a readily mobilizable pool which defines the tissue’s capacity to respond to cold challenges, highlighting their importance for metabolic homeostasis^14,15^.

Since adipose tissue remodelling coincides with the transition from maternal milk to solid food, we hypothesised that dietary changes may be a potential driver of this process. To test this, we conducted detailed phenotyping of adipose tissue taking into consideration sex dimorphism and starting at post-natal day 21, under both standard weaning conditions (pups separated from their mother at p21) and delayed weaning conditions (pups maintained with their mother until p28). We uncover sexually dimorphic mechanisms of adipose tissue expansion during the peri-weaning period, driven by a remodelling of the iWAT microenvironment.

## Results

### Post weaning remodelling of the adipose tissue is sexually dimorphic

To characterise metabolic adaptations during the weaning transition, we first measured metabolic parameters of C57BL/6J mice at p21 (weaning), p28 and p35 across sexes. The post-weaning period is associated with rapid weight gain (Figure 1A), while fat and lean mass proportions remain stable in both males and females (Figures 1B and C). This stable body composition is corroborated by the proportional increase of energy expenditure relative to body mass between p21 and p35 (Figure 1D), and unchanged levels of plasmatic leptin (Figure S1A). However, both inguinal and perigonadal WAT (gWAT) masses increase sharply during this period (Figures 1E, 1F, S1B and S1C), while interscapular BAT proportion is reduced (Figures 1G and S1I). Notably, iWAT and gWAT expansions follow different kinetics, leading to a sharp increase in the visceral index (Figure 1H). This rapid gWAT growth is accompanied by a marked increase in lipid droplet size in both sexes (Figures S1D-S1F), indicative of adipocyte hypertrophy. In contrast, the increase in iWAT proportion to total mass (Figures 1E and S1B), and iWAT adipocyte hypertrophy are only observed in p35 females (Figures 1I-1K). Changes in iWAT cellular morphology are consistent with a whitening of adipose tissue over the post-weaning period, confirmed by the downregulation of UCP1 at p35 (Figure 1L). Of note, this transient expression of UCP1 also occurs in female - but not male - gWAT (Figure S1J), suggesting a developmental predisposition to the higher gWAT beiging capacity of adult females^16^. This early sexual dimorphism is also exemplified by a higher fat mass (Figure1 B), a lower lean mass (Figure 1C), and a lower visceral index (Figure 1H) in females at p35. These findings indicate that sexually dimorphic patterns of adipose tissue distribution and function emerge during the post-weaning period.

**Figure 1.**
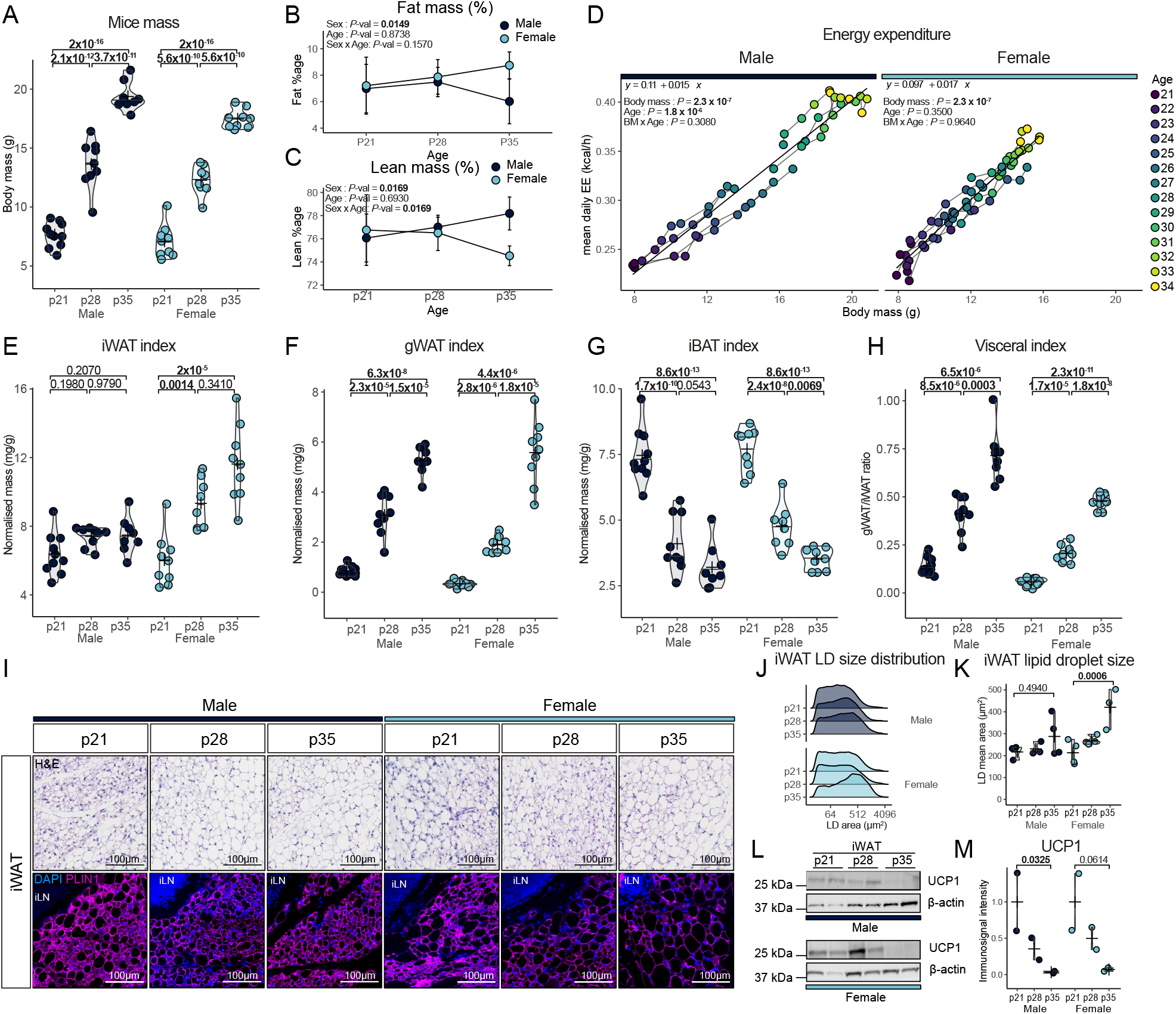
iWAT undergoes sex dimorphic remodelling post-weaning. (A) Body mass of female and male mice at weaning (p21), p28 and p35 (n = 8-10/group). (B and C) (B) Fat percentage and (C) lean mass percentage measured by magnetic resonance imaging (male: n = 8, female: n = 7). Data are represented as mean ± SD. (D) Kinetics of mean daily energy expenditure as a function of body mass from p21 to p34 (male: n = 7, female: n = 8). (E, F, G and H) Relative mass of inguinal WAT (E), perigonadal WAT (F) and interscapular BAT (G) and visceral index (H) at p21, p28 and p35 in male and female mice (n = 8-10/group). (I) Hematoxylin and eosin staining (top) and PLIN1 immunofluorescent labelling (bottom) of iWAT from p21, p28 and p35 male and female mice. (J and K) (J) Ridge plot showing the distribution of lipid droplet area and (K) mean lipid droplet area in male and female iWAT (n = 3-4/group). (L) Western blot (left) and quantification (right) of UCP1 expression in iWAT from p21, p28 and p35 male and female mice (n = 2/group). *P*-values were determined by two-sided t.test for normally distributed condition or two-sided Mann-Whitney test (A, E-H, K and L), Scheirer-Ray-Hare test followed by Dunn’s test (B and C) or ANCOVA (D)

### Microenvironment remodelling during post-weaning adipose tissue whitening

To explore the molecular mechanisms driving iWAT whitening across the post weaning period, we performed a transcriptomic analysis of iWAT at p21 and p28 in both sexes. In addition to the expected enrichment of gene sets associated with negative regulation of thermogenesis, Gene Set Enrichment Analysis (GSEA) reveals an upregulation of genes associated with immune response and innervation in both males and females between p21 and p28 (Figures 2A and 2B). This raises the possibility that, as in adult WAT browning, immune cell signalling and remodelling of the sympathetic innervation may contribute to post-weaning iWAT whitening.

**Figure 2.**
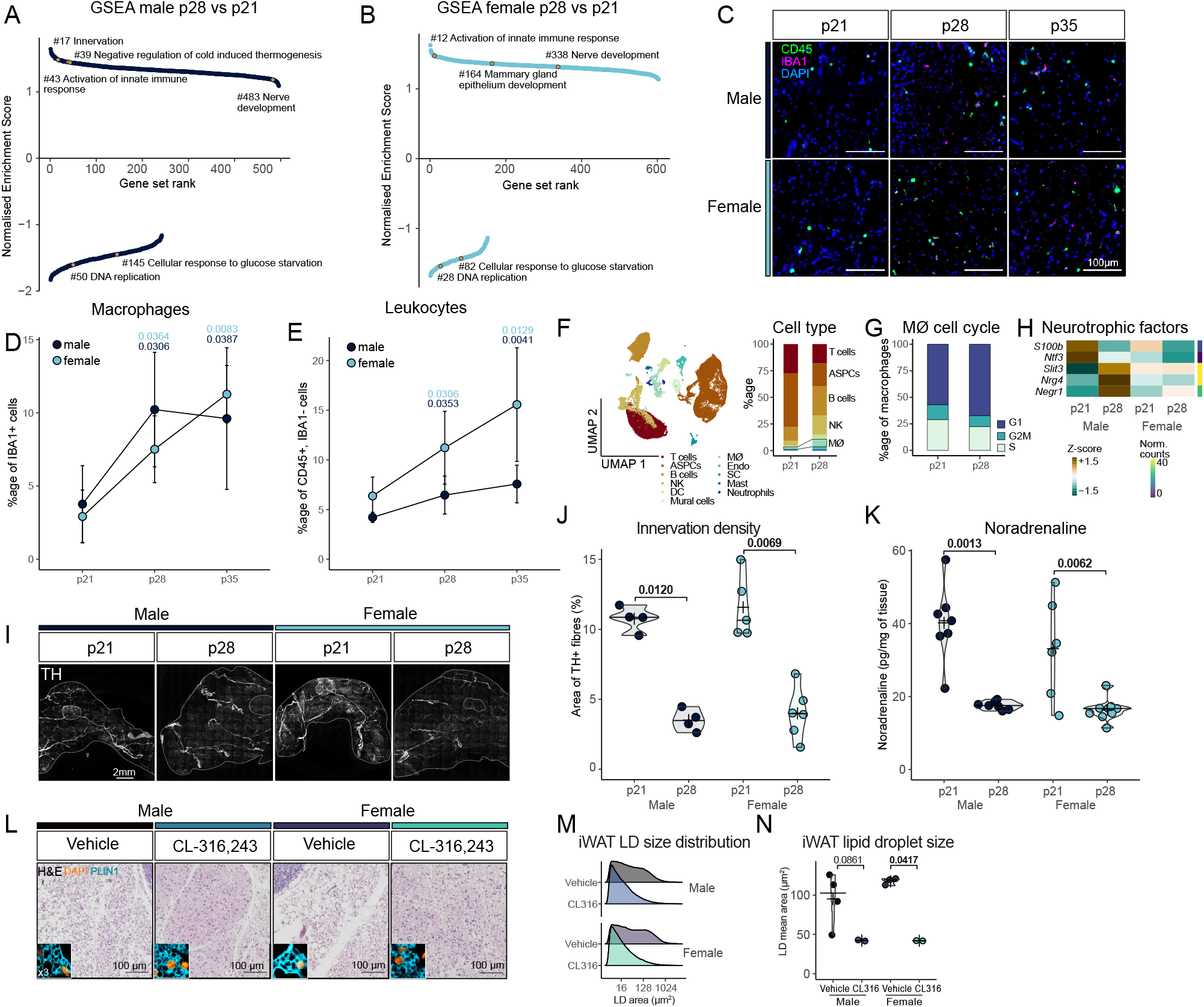
iWAT whitening associates with immune recruitment and remodelling of the innervation. (A and B) Ranked enrichment score plot from gene set enrichment analysis (GSEA) ran on MsigDB C5 GO database in DW vs SW p28 male (A) and female (B) iWAT. (C) Immunofluorescent labelling of IBA1 (magenta) and CD45 (green) of iWAT at p21, p28 and p35. (D and E) Quantification of (D) IBA1-positive and (E) CD45-positive/IBA1-negative cell proportion in p21, p28 and p35 in male and female mice (n = 2-6/group). Data are represented as mean ± SD. (F) UMAP and proportion of iWAT SVF cell type in p21 and p28 males from Qian et al single nuclei RNA-seq. (KG) Proportion of macrophages from (F) in G1, G2M or S phase. (H) Heatmap showing relative expression (z-score) andmeanabsoluteexpression(normalisedcounts) of differentially expressedgenes encoding neurotrophic factors. (I and J) (I) Whole-mount immunofluorescent analysis of TH positive innervation and (J) quantification of TH positive area in male and female iWAT at p21 and p28 (n = 4-6/group). (K) Noradrenaline content in iWAT from p21 and p28 male and female mice (n = 6-9/group). (L) Hematoxylin and eosin staining of iWAT from p21 male and female mice treated for 3 days with CL-316,243. (M and N) (M) Ridge plot showing distribution of lipid droplet area and (N) mean lipid droplet area in male and female iWAT (n = 2/group). GSEA: Gene Set Enrichment Analysis, ASPCs, adipose stem and progenitor cells, NK : natural killer cells, DC : dendritic cells, MØ : macrophages, Endo : endothelial cells, SC : Schwann cells, Mast : mast cells, SVF : stromal vascular fraction *P*-values were determined by two-sided t.test for normally distributed condition or two-sided Mann-Whitney test. (I, J and M) or Scheirer-Ray-Hare test followed by Dunn’s test (D and E)

Consistent with our transcriptomic data, co-immunofluorescence staining for the pan leukocyte marker CD45 and the macrophage marker IBA1 shows an increase in iWAT’s macrophage content, along with other leukocytes, from p21 to p35 in both male and female mice (Figures 2C-2E). To further investigate whether this increase in macrophage content reflects local proliferation or immune cell infiltration, we reanalysed an age-matched single cell RNA-seq dataset generated from the stromal vascular fraction of male mice at p21 and p28^17^. This independent dataset confirms the relative increase in iWAT immune cell content (Figure 2F) and reveals that a large subset of macrophages (>30%) are proliferating at either time point (Figure 2G), suggesting that the macrophage accumulation in iWAT is primarily driven by the proliferation of resident macrophages.

As innervation also emerged as a key pathway from our transcriptomic analysis, we next examined the expression of neurotrophic factors between p21 and p28. Males show higher expression of *Slit3, Nrg4* and *Negr1* at p28 along with reduced expression of *S100b* and *Ntf3*, while females also display decreased expression of *S100b* (Figure 2H), pointing to a remodelling of the axon network during that period. To quantify innervation levels, we performed whole-mount immunofluorescence against tyrosine hydroxylase (TH). We observe a decrease in the density of TH-positive processes (Figures 2I and 2J), and a reduction of intra tissular noradrenaline between p21 and p28 in both sexes (Figure 2K). These findings led us to hypothesise that reduced noradrenaline levels may contribute to iWAT whitening. To test this, we assessed whether pre-weaning iWAT could respond to a sympathomimetic stimulation – taking into consideration previous evidence that postnatal beiging dynamics of the iWAT are only partially influenced by innervation^7^, possibly due to limited sensitivity to noradrenaline. We administered the ß3-adrenoreceptor agonist CL-316,243 at p18, p19 and p20. This treatment leads to reduced iWAT lipid droplet size at p21 in both male and female (Figures 2J-2L), confirming iWAT ability to respond to noradrenaline via the ß3-adrenoreceptor pathway.

Together, these results highlight a coordinated remodelling of the immune and neural components in both male and female iWAT during the post-weaning period. Furthermore, they point to decreased noradrenergic signalling as a likely contributor to the iWAT whitening process during this post-natal development window.

### Prolonged breastfeeding rapidly increases adiposity in a sex-dependent manner

In natural-like conditions, mouse pups start eating solid food between p16 and p21, with the last occurrence of suckling observed around p27^18,19^. To understand the role of this dietary change on white adipose tissue remodelling, we induced delayed weaning in pups by keeping them with their mothers until p28 (delayed weaning: DW) while the rest of their litter was separated at p21 (standard weaning: SW) (Figure 3A). We first confirmed that mice kept with their mother still consume milk by examining their stomach’s content at p28, which appears lighter and more liquid in DW pups (Figure 3B). In addition, transcriptomic analyses showed increased expression of the *de novo* lipogenesis genes *Acly, Acss2, Acaca* and *Fasn* in p28 SW iWAT compared to p21, consistent with a switch to a carbohydrate-rich diet^20^, while their expression remain unchanged in p28 DW iWAT (Figure 3C).

**Figure 3.**
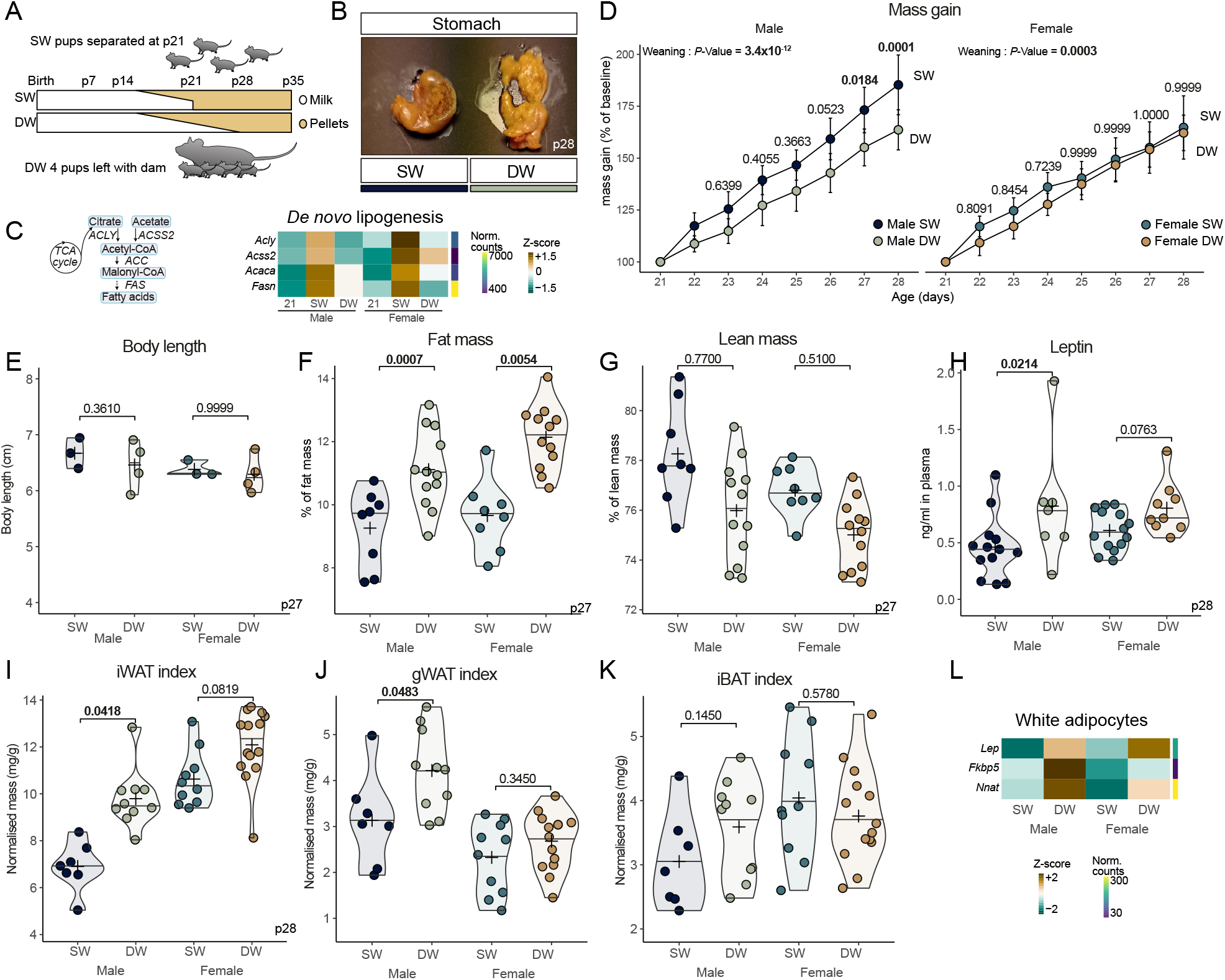
Delayed weaning promotes fat accumulation. (A) Schematic of experimental setup and expected milk to solid diet transition. (B) Stomach content of SW and DW male mice at p28 (C) Schematic of *de novo* lipogenesis (DNL) highlighting theenzymesinvolvedinthisprocess(left) and heatmap showing relative expression of DNL genes across age, sex and weaning (right). (D) Body mass gain from p21 to p35 in male and female SW and DW mice (n = 6-9/group). Data are represented as mean ± SD. (E) Body length of SW and DW male and female mice at p27 (n = 3-4/group). (F and G) (F) Fat percentage and (G) lean mass percentage as measured by magnetic resonance imaging in p27 SW and DW mice (n = 8-12/group). (H) Plasmatic leptin levels in p28 male and female SW and DW mice (n = 7-14/group). (I-K) (I) iWAT, (J) gWAT and (K) iBAT relative mass in p28 male and female SW and DW mice (n = 7-14/group). (L) Heatmap showing relative expression of white adipocytes associated genes *Lep, Fkbp5* and *Nnat*. *P*-values were determined by two-sided t.test for normally distributed condition or two-sided Mann-Whitney test (E-K) or Scheirer-Ray-Hare test followed by Dunn’s test (D)

We next examined how prolonged breastfeeding influences metabolic parameters. We observe that weaning timing has a sex-specific impact on body mass, with DW males gaining less mass than SW (Figure 3D), while body length is not affected (Figure 3E). Strikingly, both males and female DW mice show a higher proportion of fat mass at the expense of lean mass (Figure 3F and 3G). This is further corroborated by higher plasmatic concentrations of leptin in DW males compared to SW, whereas no overt difference is detected in females (Figure 3H). In line with their high adiposity, DW males have heavier iWAT and gWAT pads relative to their body mass than SW littermates, with iBAT remaining unchanged (Figures 3I-3K). A similar trend is also observed in DW females iWAT (Figure 3I-3K), despite their comparable body mass (Figure 3D). This iWAT expansion is accompanied by a robust increase in the expression of white adipocytes-related genes *Fkbp5, Nnat*^21^ and *Lep* in DW vs SW in both sexes (Figure 3L), consistent with iWAT contribution to plasmatic leptin concentration^22^. Notably, measures of food intake and energy expenditure in DW and SW mice between p29 and p35 reveal no overt difference in energy balance across conditions (Figures S2A-S2C).

Considering the critical role of steroids on adipose tissue physiology, we assessed whether DW affects gonads and adrenal glands. Steroid producing organs display similar morphology and mass between conditions at p28 (Figures S2D-S2G). The corticosteroids, corticosterone and 11-deoxycorticosterone plasmatic concentrations are not affected by timing of weaning, while the sex steroids testosterone and progesterone are barely detected in most of SW and DW samples (Figure S2H). We conclude that prolonged breastfeeding has no overt effect on steroid-producing organs.

Altogether, these findings indicate that prolonged breastfeeding promotes increased adiposity, with a more pronounced effect in males, independently of changes in steroid hormone production.

### Prolonged breastfeeding increases preadipocytes proliferation

Adipose tissue can expand either by increasing adipocyte size (hypertrophy) or number (hyperplasia)^23^. In DW females, histological analysis and PLIN1 immunostaining of iWAT both reveal larger lipid droplets compared to SW (Figures 4A-4C), pointing to hypertrophy as a primary driver of expansion. In contrast, the absence of lipid droplet enlargement in DW males suggests a role for hyperplasia.

**Figure 4.**
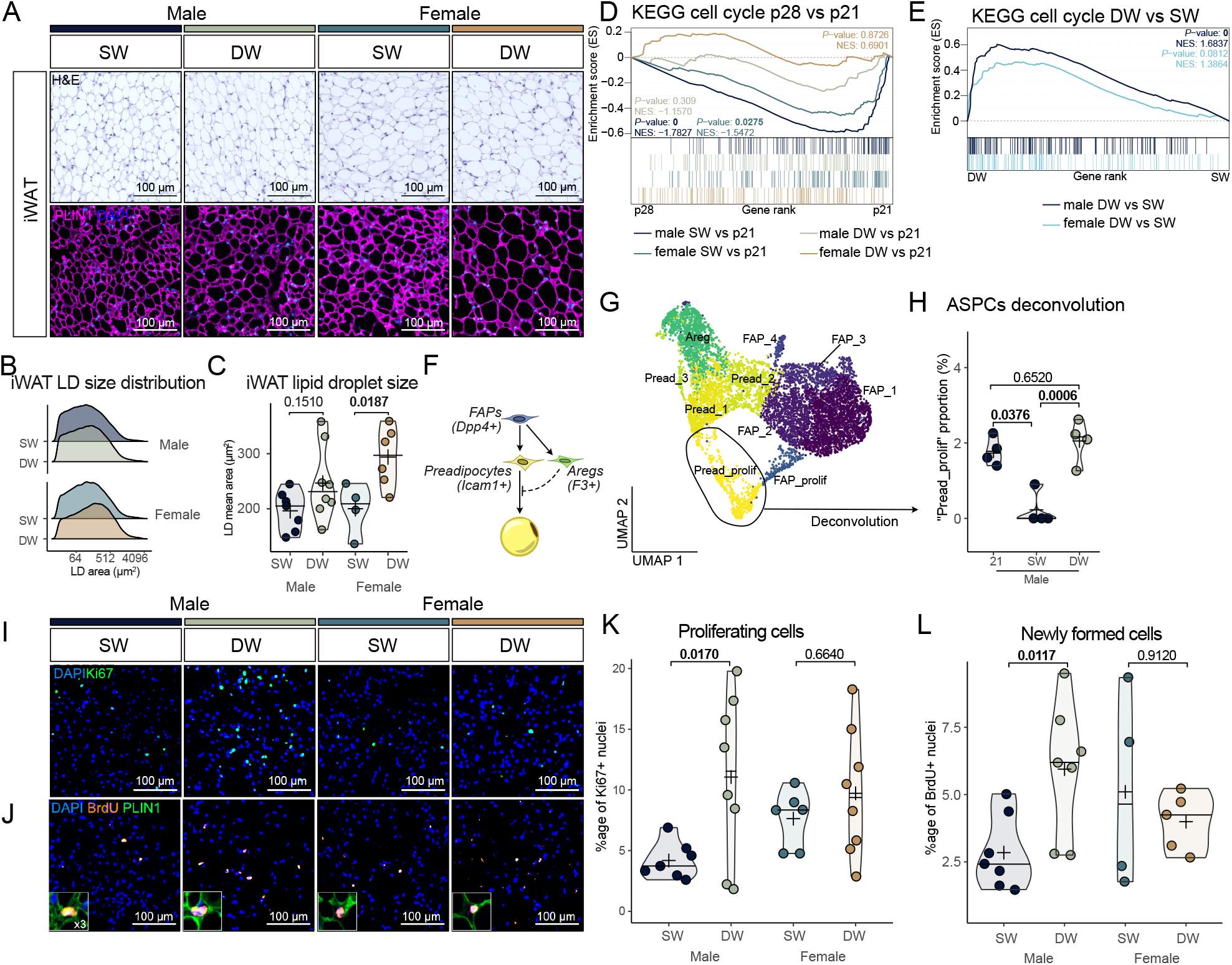
Prolonged breastfeeding promotes iWAT hypertrophy in female and hyperplasia in male. (A) Hematoxylin and eosin staining (top) and PLIN1 immunofluorescent labelling (bottom) of iWAT SW and DW male and female mice. (B and C) Ridge plot showing distribution of lipid droplet area (B) and mean lipid droplet area (C) in male and female SW and DW iWAT (n = 4-8/group). (D and E) GSEA plots of cell cycle pathway from KEGG database in (D) p28 SW and DW vs p21 iWAT and in (E) DW vs SW p28 male and female iWAT. (F) Simplified schematic of adipose tissue renewal from adipose stem and progenitor cells. (G) Reclustering of Qian et al^18^ single nuclei data set “adipose stem cells and precursors” cluster. (H) Proportion of “Pread_prolif” cells inferred from deconvolution of bulk RNA-seq data with single nuclei cell signatures derived from (I) (n=4). (I) Immunofluorescent labelling of Ki67 (green) in SW and DW iWAT at p28. (J) Immunofluorescent labelling of BrdU (orange) and PLIN1 (green)in SW and DW iWAT at p28. (K) Quantification of Ki67 positive cell proportion in SW and DW iWAT at p28 (n = 6-8/group). (L) Quantification of BrdU positive cell proportion in SW and DW iWAT at p28 (n = 4-7/group). *P*-values were determined by two-sided t.test for normally distributed condition or two-sided Mann-Whitney test.

Earlier studies have shown that proliferation of iWAT adipose stem and progenitor cells (ASPCs) gradually decreases after weaning^11,17^. This is supported by our GSEA analysis, which reveals a decrease in cell cycle-associated gene set between p21 and p28 under SW conditions (Figure 2A, 2B and 4D). In contrast, cell cycle gene expression is maintained in DW iWAT over the same period (Figures 4D and 4E), suggesting a sustained ASPC proliferation. To investigate this, we analysed ASPC proliferation in SW conditions between p21 and p28 using the same single cell RNA-seq dataset as we previously used^17^. Consistent with earlier findings, the proportion of proliferating ASPCs also decreases after weaning (Figures S3A-S3C). Reclustering of the dataset allows us to identify the 3 main ASPC subtypes^**24**^: *Dpp4+* fibroadipogenic progenitors (5 clusters), *Icam1+* preadipocytes (4 clusters) and *F3+* adipogenesis regulators (1 cluster) (Figures 4F, 4G and S3D). Deconvolution of our bulk RNA-seq data using these 10 clusters reveals no change in the proportions of the three ASPC types between p21 and p28 (Figure S3E), in line with previous reports^17^. Proliferating ASPCs are included in a *Icam1*+/*Mki67*+ cluster (“Pread_prolif”) (Figures 4G, S3B-S3D), which proportion decreases in SW iWAT between p21 and p28. Notably, this cluster is maintained in DW iWAT (Figure 4H), consistent with sustained ASPC proliferative capacity. This is supported by immuno-staining of the proliferation marker Ki67 and bromodeoxyuridine (BrdU) incorporation, both of which show increased proportion of proliferating cells in DW males compared to SW (Figures 4I-4L). In contrast, we find no increase in proliferation in female DW iWAT (Figures 4I-4L), confirming a hypertrophic rather than hyperplasic expansion.

To test whether the differences in tissue proliferation between SW and DW are cell-autonomous or driven by the microenvironment, we isolated ASPCs from DW and SW iWAT at p28 (Figure S3F), thereby eliminating environmental factors^**25**^. BrdU incorporation in 2D cultures shows a similar proliferation capacity of iWAT ASPCs between SW and DW mice (Figures S3G and S3H), suggesting that higher proliferation in DW iWAT is dependent on the tissular environment.

Our data together show that prolonged breastfeeding promotes adipose tissue growth via sex-dependant mechanisms involving a surge in progenitor proliferation in male pups, likely driven by a remodelling of adipose tissue microenvironment.

### Whitening of iWAT is potentiated by breastfeeding

Since breast milk feeding positively impacts iWAT beiging^26,27^, we hypothesised that weaning from the maternal diet could be the main driver for iWAT whitening. Unexpectedly, GSEA reveals a negative enrichment of gene sets associated with adaptive thermogenesis and oxidative phosphorylation in DW compared to SW male iWAT (Figures 5A and 5B), suggesting a loss of beige adipocytes. This is further supported by a reduced number of UCP1-positive cells, and a decrease in UCP1 protein levels (Figures 5C-5E) in males DW iWAT. While transcriptomic data suggest a similar trend for DW females (Figures 5A), we detect no changes in UCP1 protein expression (Figure 5C-5E). Thus, prolonged breastfeeding results in a sex-dependant acceleration of the iWAT developmental whitening process.

**Figure 5.**
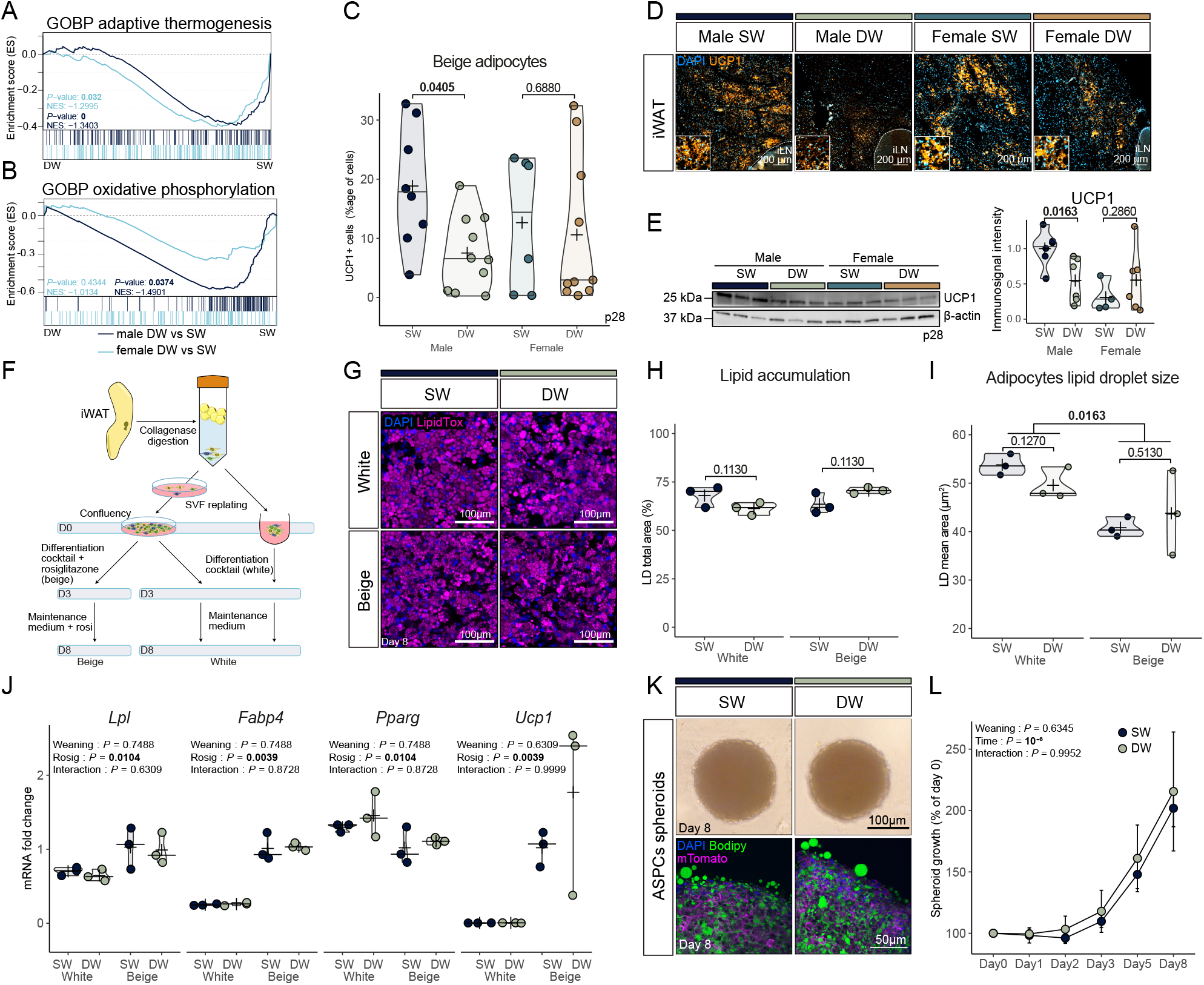
Delayed weaning accelerates iWAT whitening in a non cell autonomous fashion. (A) GSEA plot of “Adaptive thermogenesis” genes from MsigDB C5 GO database in DW vs SW p28 male and female iWAT. (B) GSEA plot of “Oxidative phosphorylation” genes from MsigDB C5 GO database in DW vs SW p28 male and female iWAT. (C and D) (D) Immunofluorescent analysis of UCP1 (orange) and (C) quantification of UCP1 positive cell proportion in p28 SW and DW iWAT (n = 6-10/group). (E) Western blot and quantification of UCP1 in p28 SW and DW iWAT protein extracts normalised to β-actin (n = 4-6/group). (F) Simplified scheme of iWAT SVF isolation and differentiation. (G-I) LipidTox (purple) labelling (G), total lipid droplet area (H) and mean lipid droplet area (I) of day 8 adipocytes differentiated from SW or DW male iWAT primary ASPCs using a white or differentiation protocol (n = 3/group). (J) Gene expression of differentiation markers *Lpl, Fabp4, Pparg* and *Ucp1* as determined by qPCR (n = 3/group). (K) Brightfield (top) and fluorescent staining (bottom) of spheroids grown from SVF isolated from SW or DW male iWAT. Nuclei are stained by DAPI (blue), lipid droplets by bodipy (green) and cells express a membrane associated myristoylated form of tomato (purple). (L) Quantification of spheroids growth based on brightfield pictures (n = 4/group).Data are represented as mean ± SD. SVF : stromal vascular fraction, ASPCs : Adipose Stem and Progenitor Cells *P*-values were determined by two-sided t.test for normally distributed condition or two-sided Mann-Whitney test (C, E,H,I) or Scheirer-Ray-Hare test followed by Dunn’s test (J and L)

Prolonged breastfeeding promotes hyperplasic iWAT expansion in males, as opposed to hypertrophic growth in females (see Figure 4), suggesting that male-specific iWAT whitening in DW is driven by increased cellularity due to *de novo* white adipocyte differentiation. Since ASPC differentiation capacity decreases after weaning^28^, we reasoned that prolonged breastfeeding may reprogram ASPCs towards an increased adipogenic potential and/or a decreased beiging ability. To test this, we performed *in vitro* white and beige differentiation using ASPCs isolated from male SW and DW iWAT (Figure 5F). Staining of neutral lipid with LipidTox shows that both induction cocktails result in a comparable differentiation efficiency of ∼70 % (Figures 5G and 5H), with no difference in mean lipid droplet size (Figures 5G and 5I) or in the expression of key adipogenic and thermogenic markers (*Lpl, Fabp4, Pparg* and *Ucp1)* between SW and DW (Figure 5J). Similarly, 3D adipose spheroid cultures from both conditions show similar growth and lipid accumulation (Figures 5K and 5L). These results indicate that prolonged breastfeeding does not intrinsically alter ASPC differentiation and imply that the reduced proportion of beige adipocytes in male DW iWAT is likely driven by endocrine or paracrine factors.

### Adipocytes in iWAT recapture noradrenaline upon prolonged breastfeeding

Having established that iWAT is responsive to noradrenaline via the ß3-adrenoreceptor pathway at weaning (Figure 2) and given that increased leptin production (Figure 3), increased ASPC proliferation (Figure 4) and reduced beige adipocyte proportion (Figure 5) are hallmarks of decreased sympathetic tone^29–33^, we next examined adipose tissue innervation. We find that prolonged breastfeeding induces a marked, male-specific increase in iWAT innervation. Transcriptomic analysis reveals that only DW males exhibit an enrichment of genes associated with the term “regulation of neuron projection development” (Figure 6A). Consistently, we detect an increase in mRNA levels of the pan neuronal marker *Ncam1* in DW compared to SW male iWAT, while no difference is observed in females (Figure S4A). Whole mount immunostaining of TH in iWAT further confirms this increase in the relative innervation of male DW mice’s iWAT depots (Figures 6B and 6C), although total TH protein levels remain unchanged (Figure S4B).

**Figure 6:**
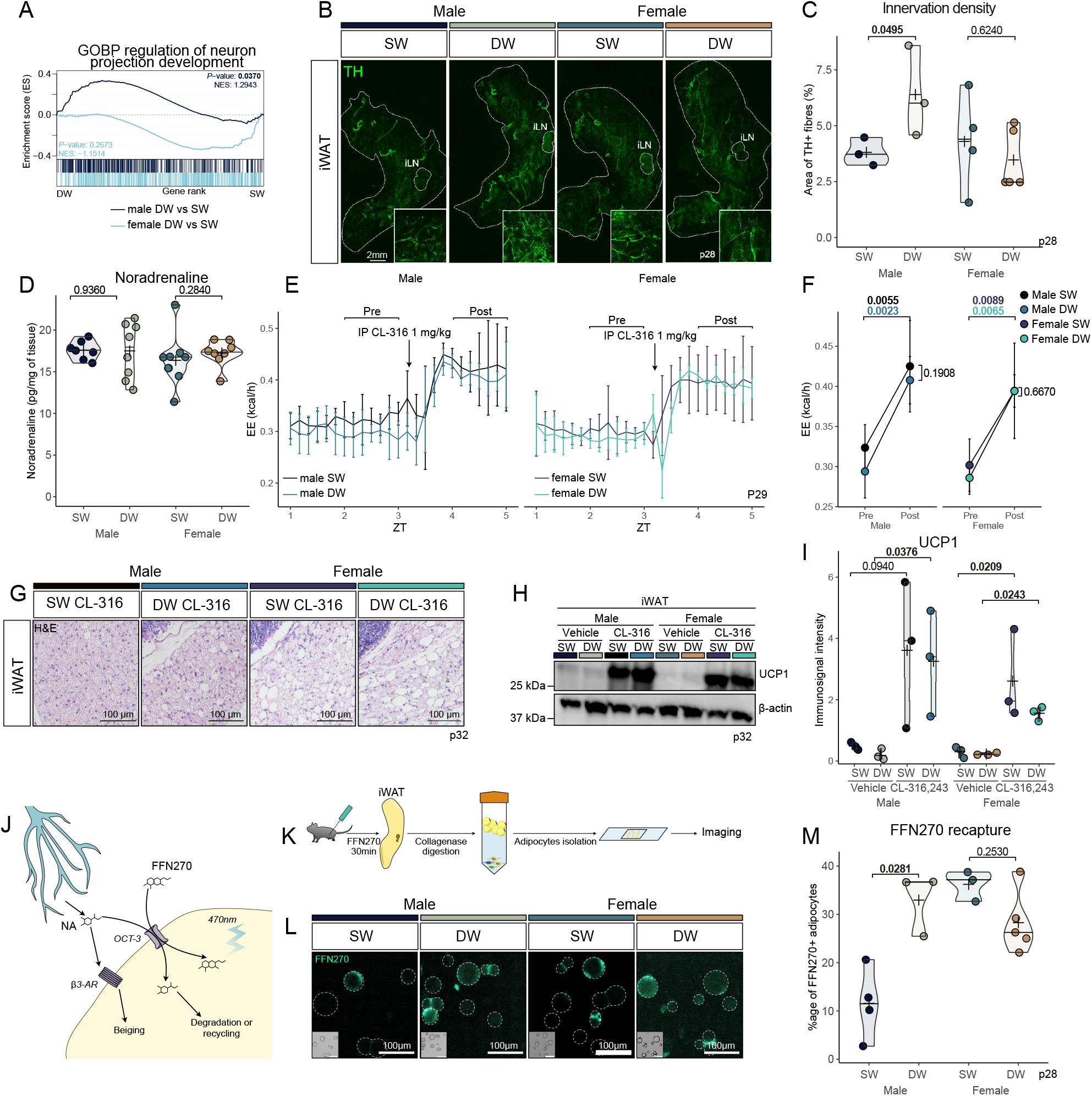
Noradrenaline is inactivated through adipocytes uptake. (A) GSEA plot of “Regulation Of Neuron Projection Development” genes extracted from MsigDB C5 GO database in DW vs SW p28 male and female iWAT. (B and C) (B) Whole-mount immunofluorescent analysis of TH positive innervation and (C) quantification of TH positive area in p28 SW and DW iWAT (n = 3-5/group). (D) Noradrenaline content in iWAT from SW and DW p28 mice (n = 7-8/group). (E and F) Energy expenditure before and after CL-316-243 i.p. injection (0.1mg/kg) (n = 4-5/group). Data are represented as mean ± SD. (G) Hematoxyllin and eosin staining of p32 iWAT from SW and DW male and female mice treated for 3 days with CL-316,243. (H and I) (H) Western blot analysis and (I) quantification of UCP1 normalised to β-actin in p32 iWAT of SW and DW mice treated with CL-316,243 or PBS for 3 days (n = 3/group). (J) Schematic illustrating noradrenaline and FFN270 uptake by adipocytes. (K) Protocol for FFN270 *in vivo* labelling followed adipocytes isolation and imaging. (L and M) FFN270 labelling with brightfield insets of isolated adipocytes from p28 SW and DW male and female iWAT (left) and quantification of the proportion of adipocytes labelled with FFN270 after 30 minutes incubation (right) (n = 3-5/group). *P*-values were determined by two-sided t.test for normally distributed condition or two-sided Mann-Whitney test (C, D, F, and M) or Scheirer-Ray-Hare test followed by Dunn’s test (I)

To reconcile the observed iWAT whitening phenotype with the increased innervation in DW mice, we considered three possible mechanisms: (i) a decrease in the tissue’s noradrenaline content, (ii) a post synaptic resistance to noradrenaline, or (iii) an enhanced noradrenaline reuptake. We first measured the total noradrenaline content in the iWAT, which is similar for all conditions at p28 (Figure 6D). Next, we assessed adipocytes sensitivity to noradrenaline by treating p29 DW and SW mice with the ß3-adrenoreceptor agonist CL-316,243. Acute treatment leads to a similar increase in energy expenditure in both SW and DW mice (Figures 6E and 6F), while a three-day treatment induces iWAT beiging and UCP1 protein accumulation in all conditions (Figures 6G-6I).

Having ruled out both reduced noradrenalin content and impaired adipose tissue responsiveness, we next considered alterations in noradrenaline clearance. Noradrenaline can be recaptured by sympathetic neuron-associated macrophages (SAMs) through the transporter SLC6A2^34^ or by adipocytes through SLC22A3 (OCT-3)^35,36^ before degradation by the Monoamine Oxidase A (MAO-A)^34,37^ (Figure 6J). Notably, *Maoa* expression is increased in male DW iWAT vs SW (Figure S4C), supporting the hypothesis of enhanced noradrenaline clearance. However, only a small subset (6%) of stromal vascular fraction (SVF) cells express *Maoa* in p21 or p28 scRNA-seq data (Figure S4D), and none of these co-express noradrenaline transporters (Figure S4D). Deconvolution of immune cell populations from our bulk RNA-seq data reveals no difference in immune cell composition between DW and SW (Figure S4E). Furthermore, macrophage marker expression does not indicate increased macrophages abundance (Figure S4F). Finally, SAM-specific markers are not differentially expressed between SW and DW iWAT, ruling out an expansion of this macrophage subpopulation (Figure S4G). In contrast to other monoamines transporters, the adipocyte-specific noradrenaline transporter *Slc22a3* is readily detected in our iWAT bulk RNAseq dataset (Figure S4H), pointing to adipocytes as likely contributors to noradrenaline reuptake.

To test the capacity of SW and DW adipocytes to recapture noradrenaline, we injected p28 mice transcutaneously in the inguinal fat pads with FFN270, a fluorescent, pharmacologically inert noradrenaline analogue, and isolated adipocytes from the SVF after 30 minutes^38^ (Figure 6J and 6K). As expected, FFN270 staining is absent in SVF cells (Figure S4I) confirming adipocytes as the main drivers of recapture, while a three-fold increase in FFN270-positive adipocytes is observed in male DW compared to SW iWAT, and no difference is detected in females (Figure 6L).

Altogether, these findings indicate that in males, DW adipocytes retain their responsiveness to ß-adrenergic stimulation but actively recapture noradrenaline. This enhanced clearance may explain the male-specific reduction in beige adipocytes content in DW iWAT, despite an increased innervation.

## Discussion

Our study thoroughly maps the morphological, physiological and transcriptional changes taking place in the inguinal subcutaneous adipose tissue after postnatal day 21, during the critical peri-weaning developmental window in mice. We reveal that prolonged breastfeeding elicits a sex-specific remodelling of adipose tissue, marked in males by a maintenance of tissue proliferation and an acceleration of the whitening process.

We found that the redistribution of the adipose tissue following weaning reported by previous studies^6–8,10,15^ is associated with a global whitening of the iWAT in a sexually dimorphic manner. This progressive replacement of beige adipocytes by white adipocytes has been shown to occur in adult iWAT after termination of cold exposure^21^, denervation^39^ or high fat diet^40^, mainly through the transdifferentiation of mature adipocytes^12,41,42^. Here, we demonstrate that the peri-weaning whitening is associated with a decrease in innervation density and noradrenaline concentration (Figure 2). While beige adipocytes biogenesis in iWAT before weaning occurs independently of innervation and housing temperature^7^, our data show that iWAT from p21 and p28 mice is already responsive to sympathomimetic stimuli (Figures 2 and 5), suggesting that sympathetic tone could contribute to beige adipocyte maintenance as early as p21. Prolonged breastfeeding promotes higher innervation in males, which may be mediated by increased leptin secretion (Figure 3). Indeed, leptin has been shown to exerts a positive effect on development^8^, maintenance^43^ and function^44,45^ of sympathetic nerves in the iWAT. Of note, increased leptin levels are also observed in human after prolonged breastfeeding^46^.

We observe that the higher innervation in DW iWAT is paradoxically associated with reduced beige adipocyte content (Figures 5 and 6), suggesting a post-synaptic compensation. Indeed, noradrenaline levels remain stable despite higher innervation, which may reflect increased degradation. We show that adipocytes in p28 iWAT have the capacity to recapture synaptic noradrenaline, in line with previous studies in adult mice^36,47^. This reuptake, combined with increased MAOA expression, may serve as a compensatory mechanism to fine-tune local noradrenaline concentrations. As a matter of fact, MAO-A is not expressed in adult healthy mouse iWAT^37,48^, but cold exposure in UCP1-KO mice leads to the appearance of MAO-A positive cells^37^. This suggests that MAO-A expression may be upregulated in response to noradrenaline overflow and could explain its expression during postnatal development. OCT-3, on the other hand, is expressed in a high proportion of adipocytes in mice and humans^35,36,48,49^ and recaptures noradrenaline in a concentration-dependent manner^50^. This implies that adipocytes not expressing MAO-A could still recapture noradrenaline and either release it back to the extracellular milieu or store it in vesicles^51^.

Adipose stem and progenitor cell (ASPC) proliferation normally decreases between p21 and p28^11,17^. Yet, we show here that this is prevented by prolonged breastfeeding in males (Figure 4). During this period, adipogenesis regulator (Areg) cells have been reported to switch from a proto an anti-adipogenic role^28^. This transition could be regulated by milk intake and control preadipocyte proliferation. However, our *ex vivo* assessment of ASPC proliferation and differentiation indicates that the difference in proliferation between SW and DW iWAT is caused by external factors, arguing against a major role of Areg cells. Alternatively, milk compounds could directly promote proliferation. For instance, Meln et al^52^ have shown that dietary fat such as palmitoleic acid can promote ASPC proliferation in post weaning mice independently of caloric load. Since milk contains palmitoleic acid^53,54^, this may contribute to the enhanced proliferation observed in DW iWAT.

We find that macrophage accumulation is a feature of age-related, but not weaning-induced, iWAT whitening (Figure S6). Given that macrophages can both promote^55–57^ and repress^34^ beiging, they are likely to regulate postnatal adipose remodelling. In fact, macrophages were shown to act as mediator of postnatal iWAT beiging by converting breast milk alkylglycerols into platelet activating factor^27^. In contrast, our data show that macrophage content increases with age and associates with reduced beiging, independently of breastfeeding duration (Figure 5). This discrepancy may be explained by decreased alkylglycerol content with time, as milk composition changes throughout lactation^53,54^. An alternative explaination is that milk intake after p21 may not be sufficient to counteract adipocyte-mediated degradation of alkylglycerols ^27^.

An unexpected result of our study is that we observe a strong sexual dimorphism in the phenotype elicited by prolonged breastfeeding: males display higher proliferation, innervation, and noradrenaline reuptake, while females show adipocyte hypertrophy. This dimorphism is unlikely to result from variations in sexual hormones, as mice are still prepubertal with plasmatic levels of progesterone and testosterone mostly below the detection limit (Figure S2), and the steroid profile in the first three weeks of life is not sexually dimorphic^58^. Instead, chromosomal difference may underlie this dimorphism^59^, as supported by sex differences in body composition, adipose tissue distribution and gWAT beiging already present at p21 (Figures 1 and S1). Of note, the female-specific beiging of gWAT at p21 (Figure S1) aligns with previous reports of higher beiging potential in female gWAT compared to males at 6 weeks of age^16^. In addition, female iWAT mammary glands undergo extensive remodelling between p21 and p28 (Figure 2B), involving the recruitment of PDGFRα+ progenitors^60^ and possibly adipocyte dedifferentiation, as seen during lactation^61,62^. This process was recently shown to alter beige adipocytes physiology^63^ and may contribute to the observed sex-specific differences in whitening kinetics.

Overall, our study has implication for human health, in line with numerous longitudinal studies showing beneficial effects on metabolism of children breastfed for a longer period^1,64,65^ and of milk over formula^66^ as well as long term protection from diet induced obesity offered by delayed weaning in rats^26^. It also adds up to the list of parameters to control in metabolic studies^67^, where timing of weaning should be considered and reported as a confounding factor.

## Limitation of the study

Milk is a complex liquid containing a wide array of bioactive compounds^68^, and further studies will be required to identify and isolate factors modulating the offspring’s adipose tissue maturation and function during and after lactation. In addition, while our data confirm that DW mice are still consuming milk, it remains challenging to disentangle the respective contributions of dietary transition and maternal separation. Indeed, mice gradually transition away from their mother from p14^18,19^. Furthemore, the time spent with the mother at p26 differs between male and female with male being more prone to explore^69^, potentially contributing to the observed sexual dimorphism.

## Resources availability

Raw and processed data have been deposited in NCBI’s GEO database (GSE306420). Single cell data from male iWAT ontogeny from Qian et al^17^, was downloaded from GSE249814.

## Acknowledgements

This paper was typeset with the bioRxiv word template by @Chrelli: www.github.com/chrelli/bioRxiv-word-template We would like to thank Catherine Hayward for assistance with histological tissue preparation, Hong Qu for scanning of tissue sections, Thanh Thai Pham, Sigrun Lislien and Kristine Stamrud Beich for animal care. We also thank the MolMed Imaging Platform (MIP) at the Institute of Basic Medical Sciences, University of Oslo for providing access to and training on relevant microscopes. This work was supported by the Norwegian Research Council (324281). We acknowledge the Norwegian Sequencing Centre (Oslo University Hospital) for professional sequencing services.

## Author contributions

D.D., P.C., C.B. and N.B. designed research.

K.T.D. provided access and assisted with use of metabolic cages.

D.D., S.K.S.B., N.F., F.D. and I.P. performed experiments.

D.D. analysed the data.

D.D. and N.B. wrote the paper.

D.D., S.K.S.B., F.D., I.P., K.T.D., P.C., C.B. and N.B. edited the paper.

## STAR Methods

### Key resources table

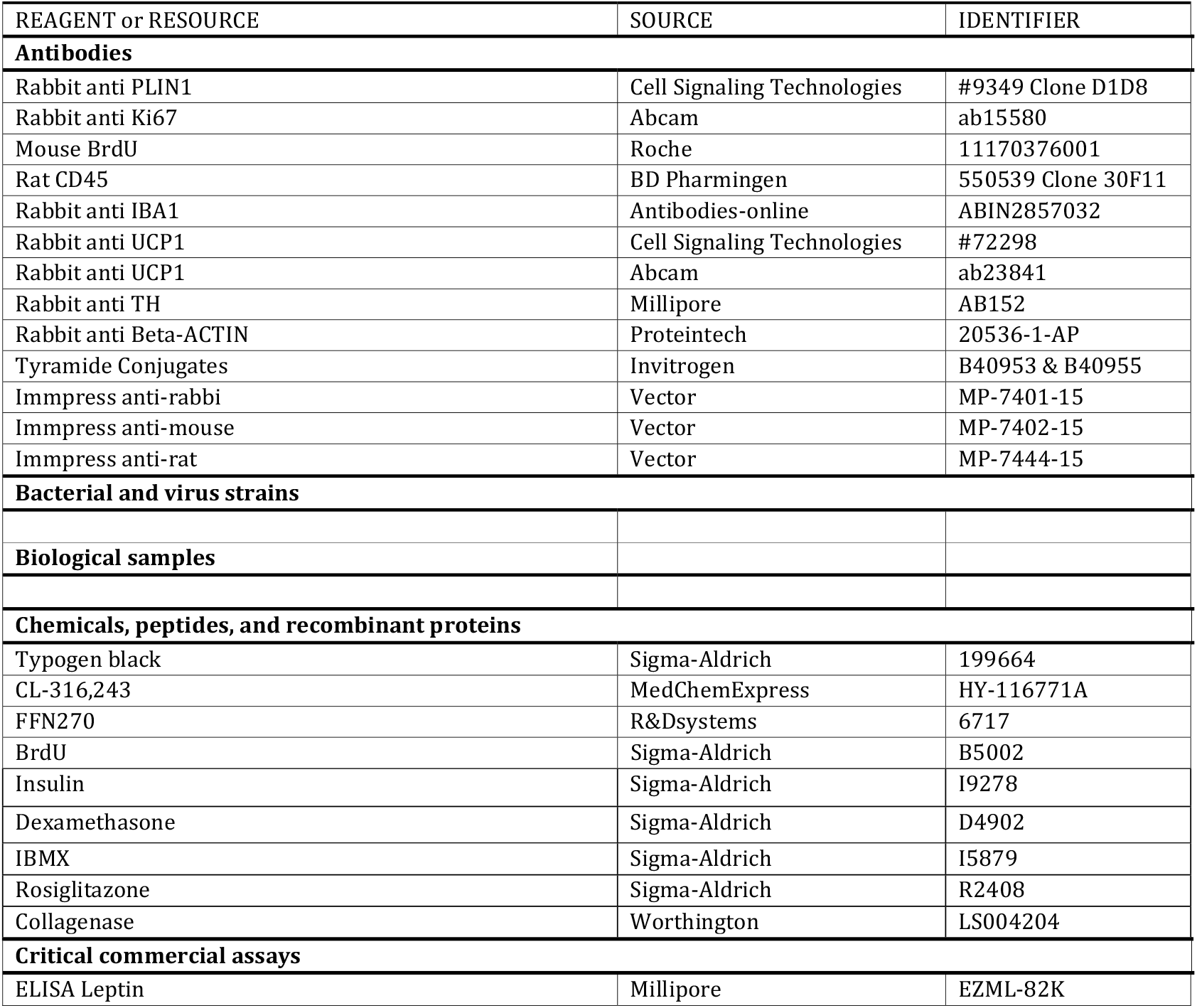
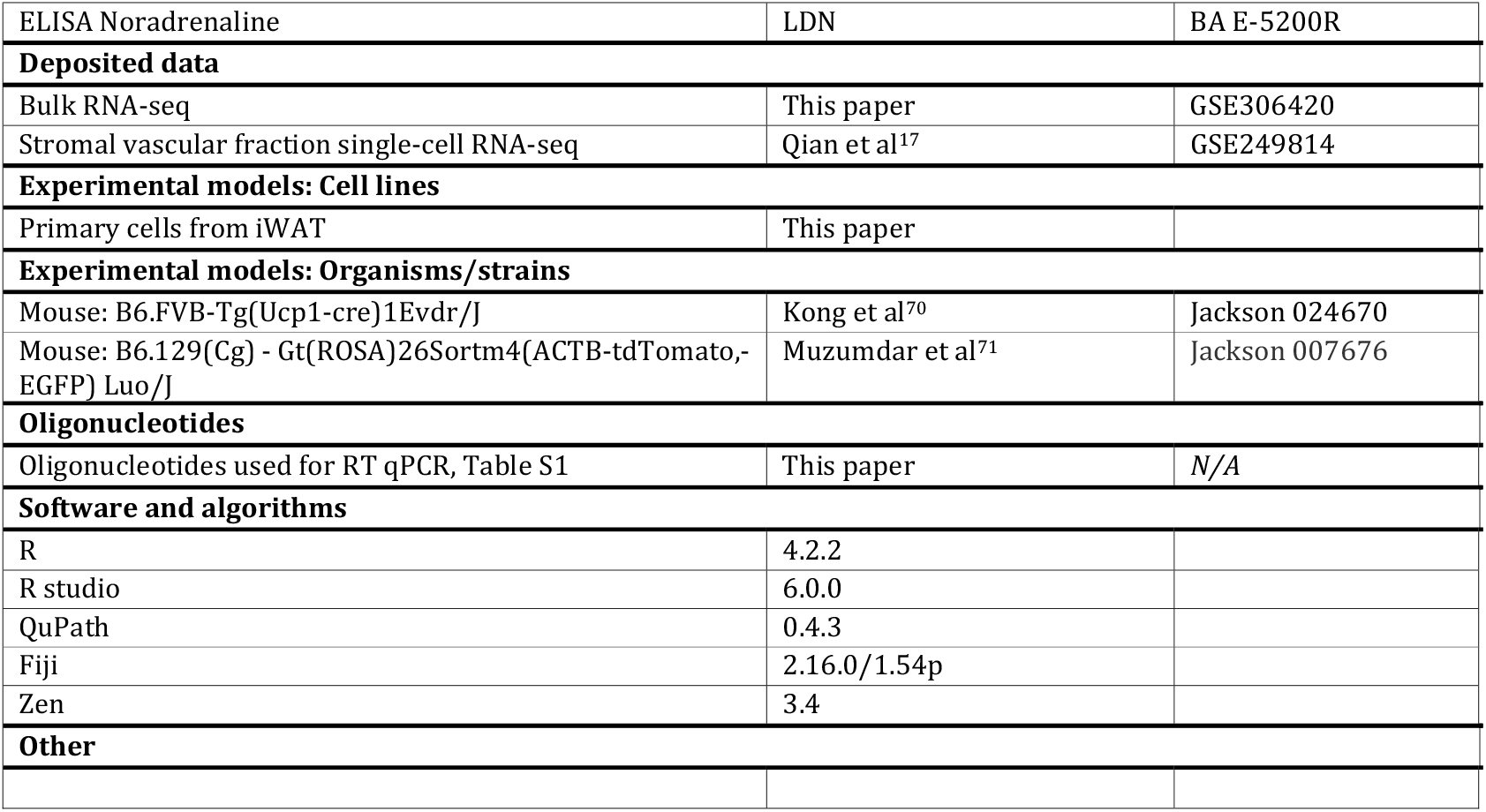

### Experimental model and subject details

#### Mice

Mice bred in-house and maintained on a C57Bl/6 genetic background were housed on a 12-h light/12-h dark cycle (lights on at 7:00 am) at room temperature. Mice were fed normal, commercial rodent chow and provided with water *ad libitum*. All mice had access to hiding houses, tunnel and paper. Unless otherwise stated, mice were sexed at p21, by approximation of anogenital distance and genital morphology, weaned and kept with same sex siblings. Mice from 6 to 10 pups’ litters were used in the study.

#### Delayed weaning protocol

Mice in the delayed weaning condition were kept in group of four with their mother to reduce variability. They were maintained in large cages (GR900) with food only available in basket attached to the lid, reducing the accessibility of chow over milk. Separated mice were in group of two to four in regular housing cages (GM500) with food pellets available in the basket and at the bottom of the cage. Littermates were systematically used as controls.

### Method details

#### Indirect calorimetry and body composition

Body composition was determined with a Minispec LF90II magnetic resonance imaging machine (Bruker, Billerica, MA, USA). Mice were single housed in an indirect calorimetry system consisting of 20 individual cages (Phenomaster, TSE Systems, Germany) and maintained at room temperature with *ad libitum* access to food. Gas flow was 0.42 L/min, with gas exchange measured for 10 seconds per cage in 20 minutes intervals. Physical activity was recorded as movements in the XY-plane. Food intake and body mass changes were measured every 10 minutes for each mouse. To measure response to sympathomimetic *stimuli*, mice were treated with CL-316,243 (0.1mg/kg) diluted in PBS and immediately replaced in the metabolic cages.

#### Immunofluorescence

Tissues were fixed in 4% PFA for 4-6h and embedded in paraffin. 5-μm sections were deparaffinised and processed for H&E. For immunohistochemistry or immunofluorescence, deparaffinised slides were submerged in antigen retrieval buffer and microwaved for 8 min. After being rinsed with PBS, slides were blocked for an hour with 2.5% horse serum (Vector) and incubated overnight at 4 °C with primary antibody. After rinsing, slides were incubated with ImmPRESS polymer for 30 min at room temperature. HRP activity was detected with Alexafluor (Thermo Fisher).

Primary cells were grown on coverslips, fixed in 4% PFA for 10 minutes then permeabilised with Triton 0.1x for 15 minutes. After blocking for 1 hour with 5% BSA in PBS, cells were incubated with primary antibodies overnight at 4C and Alexafluor coupled secondary antibody for one hour at room temperature.

Images were acquired with Zeiss Axioscan Z1 or Zeiss Observer and ZEN 3.4 blue edition software and analysed with QuPath 0.4.3 software or ImageJ.

To assess iWAT proliferation, mice were injected with BrdU (50mg/kg) 24 hours before termination.

#### Western blot

Protein lysates were extracted using Precellys tissue lyser (Bertin, France) and lysis buffer (TRIS 50 mM, Sucrose 0.27 M, EDTA 1 mM, EGTA 1mM, Na pyrophosphate 5mM, Na β-glycerophosphate 10 mM, Na orthovanate 1 mM, NaF 50 mM, triton 1%) supplemented extemporaneously with β-mercaptoethanol 0,1% and protease inhibitors (Roche, Basel, Switzerland). After blocking for 1h in 5% BSA, membranes were blotted with the primary antibodies overnight at 4°C and incubated in HRP-coupled secondary antibodies for 1h at room temperature. Images were acquired using ChemiDoc MP Imaging System camera system (Bio-Rad) and quantified using imageJ.

#### Whole mount staining

Tissues were fixed in 2% PFA for 4-6h and processed according to Willows et al^72^. Tissues were Z-depth reduced and subsequently blocked in 2.5% BSA, 1% Triton X-100 in PBS for 24h at 4°C and incubated in 0.1% Typogen Black (Sigma, Cat#199664) to reduce autofluorescence. Tissues were then incubated with primary antibodies for 48h at 4°C and secondary overnight. Stacked images (5x15µm) were acquired with Andor Dragonfly and analysed with ImageJ and QuPath 0.4.3.

#### Evaluation of noradrenaline uptake

Mice were transcutaneously injected with FFN270 (80 µl of 1 mM solution) into the iWAT. After 30 min, adipose tissue was collected, minced and incubated in 0.2% collagenase for 45 min at 37 °C. After filtering and centrifugating at 500g for 5 min at room temperature, floating adipocytes or pelleted stromal vascular fraction were resuspended in 250 µl of HBSS. 20 µl of cells solution was mounted between slide and coverslip and imaged with a Zeiss Axio Observer at 10x magnification.

#### Hormone analysis

At the end of experimental procedures mice were terminated by decapitation between 9 and 11am and trunk blood was collected in vacuum blood collection tubes (VF-053STK, Terumo). Serum was separated by centrifuging blood at 2500 rpm for 20 min at 4C. Leptin was determined by ELISA (Mouse leptin ELISA kit, Millipore), steroids were measured by LC-MSMS ^73^.

Intratissular noradrenaline was determined by homogenising the tissue in 750 µl of 0.01 N HCl, EDTA 1 mM. Cellular debris were pelleted by centrifugating at 12000rpm for 12min at 4C. Supernatant was processed according to manufacturer instructions for ELISA determination (noradrenaline high sensitive ELISA kit BA E-5200R, LDN, Germany). Values were normalised to raw mass of the tissues.

#### RNAseq

Gene expression profile was analysed for subcutaneous inguinal white adipose tissue from four p21 mice, four delayed-weaning p28 mice and four standard-weaning p28 mice of each sex. RNA extraction was performed using RNeasy Lipid Tissue micro-Kit (Qiagen) following manufacturer instructions.

Sequencing libraries were prepared using the KAPA mRNA HyperPrep kit (Roche). Sequencing was performed on NovaSeqXPlus sequencer (Illumina) with a 25B flow cell using NovaSeqXPlus System Suite: 1.2.0.28691 and BCL Convert: 4.1.23. Reads were filtered with fastp, aligned to the mm10 genome with hisat2 and counted using featureCounts.

Differential expression was performed using the DESeq2 package version 1.38.3. Gene Set Enrichment Analysis was performed using the GSEA software^74^ and plotted using an adaptation of the replotGSEA function from the Rtoolbox package (https://github.com/PeeperLab/Rtoolbox). Gene ontology analysis was performed using g:Profiler^75^. Data visualisation was carried out using R software (v4.2.2), Pheatmap package was used for heatmaps, Vennerable for Euler diagrams and ggplot2 for plots.

#### SVF single cell RNA-seq analysis

Single cell data from male iWAT ontogeny from Qian et al^17^ was reanalysed using the Seurat package with the same parameters as the initial study. 21757 features for 15365 cells defined as preadipocytes were reclustered with a Louvain algorithm, 10 principal components, and a resolution of 0.5. 10 clusters were classified by their expression of the markers *Dpp4, F3* and *Icam1* and their expression of proliferation markers *Mki67* and *Cdk1*.

Matrices from p21 and p28 preadipocytes, immune cells and all cells were generated to produce signature files which were used in CiberSortX for deconvolution of bulk RNAseq data.

#### SVF isolation

Adipose tissue from p28 SW and DW males was collected, minced and incubated in 0.2% collagenase diluted in DMEM/F12 for 45 min at 37 °C. After centrifugating at 500g for 5 min at room temperature and successive filtering on 100 µm and 40 µm strainers, stromal vascular fraction cells were plated and cultivated in basal medium (DMEM/F12 17.5 mM glucose with glutamax, 10% fetal calf serum and 1% penicillin/streptomycin). Upon confluency, adipogenic differentiation was induced by cultivating cells in basal medium supplemented with 1 µg/ml insulin, 1 µM dexamethasone and 0.5 µM IBMX, supplemented with 1 µM rosiglitazone for beige differentiation. After 3 days, medium was changed to basal medium supplemented with 1 µg/ml insulin. Spheroids were formed by seeding 50000 cells in low binding 96 well plates and induced to the following day as described above.

Cell proliferation was assessed by incubating proliferating stromal vascular fraction cells with 10µM BrdU for 6 hours and processed for immunocytofluorescence staining.

#### Statistics

Statistics were conducted using R language and Comp3Moy function from sumo package (https://github.com/Damien-Dufour/sumo). Normality of populations distribution was assessed with Shapiro & Wilk test for n ∈ [7,5000] or otherwise Kolmogorov–Smirnov normality test. If data followed a normal distribution, homoscedasticity was estimated with a Barlett test. To compare two populations, unpaired, two-tailed t test was used for normally distributed data with the same variance, Mann–Whitney for non-normal distributions and Welch t test for normally distributed data but with different variances. To compare three or more distributions: one-way ANOVA for normally distributed samples with pairwise multiple t tests or Kruskal–Wallis for nonnormally distributed samples with planned comparisons using Dunn’s test to determine the genotype effect or the treatment effect. Two-way ANOVA or Scheirer-Ray-Hare test were used to evaluate the effect of more than one factor. *P-*values in figures 1.D, S2B and S2D were determined by ANCOVA. Crosses on the violin plots represent the mean while the median is represented by a horizontal line. Error bars in barplot represent the SD unless otherwise stated. The number of samples per condition is indicated in the figure legends. Immunostaining pictures are representative of a group of at least three replicates.

---

**Figure S1.**
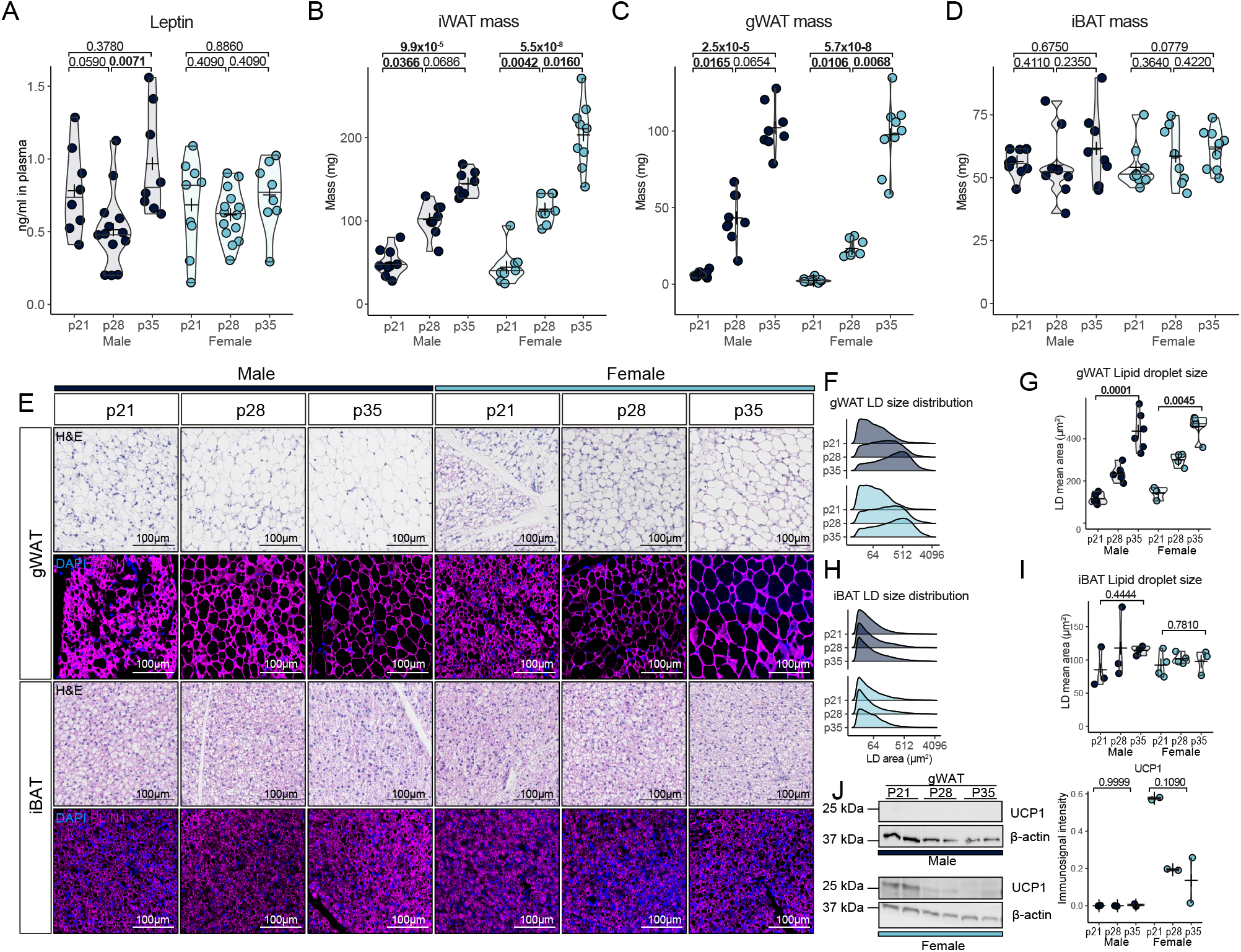
Related to Figure 1. (A) Evolution of plasmatic leptin levels in male and female (n = 8-14/group). (B,C and D) Relative mass of inguinal WAT (B) perigonadal WAT (C) and interscapular BAT (D) at p21, p28 and p35 in male and female mice (n = 8-10/group). (E) Hematoxylin and eosin staining (top) and PLIN1 immunofluorescent labelling (bottom) of gWAT and iBAT from p21, p28 and p35 male and female mice. (Fand G) (G) Ridge plot showing distribution of lipid droplet area and (G) mean lipid droplet area in male and female gWAT (n = 3-6/group). (H and I) (H) Ridge plot showing distribution of lipid droplet area and (I) mean lipid droplet area in male and female iBAT (n = 3-4/group). (J) Western blot analysis and quantification of UCP1 in gWAT from p21, p28 and p35 male and female mice (n = 2/group). *P*-values were determined by two-sided t.test for normally distributed condition or two-sided Mann-Whitney test.

**Figure S2.**
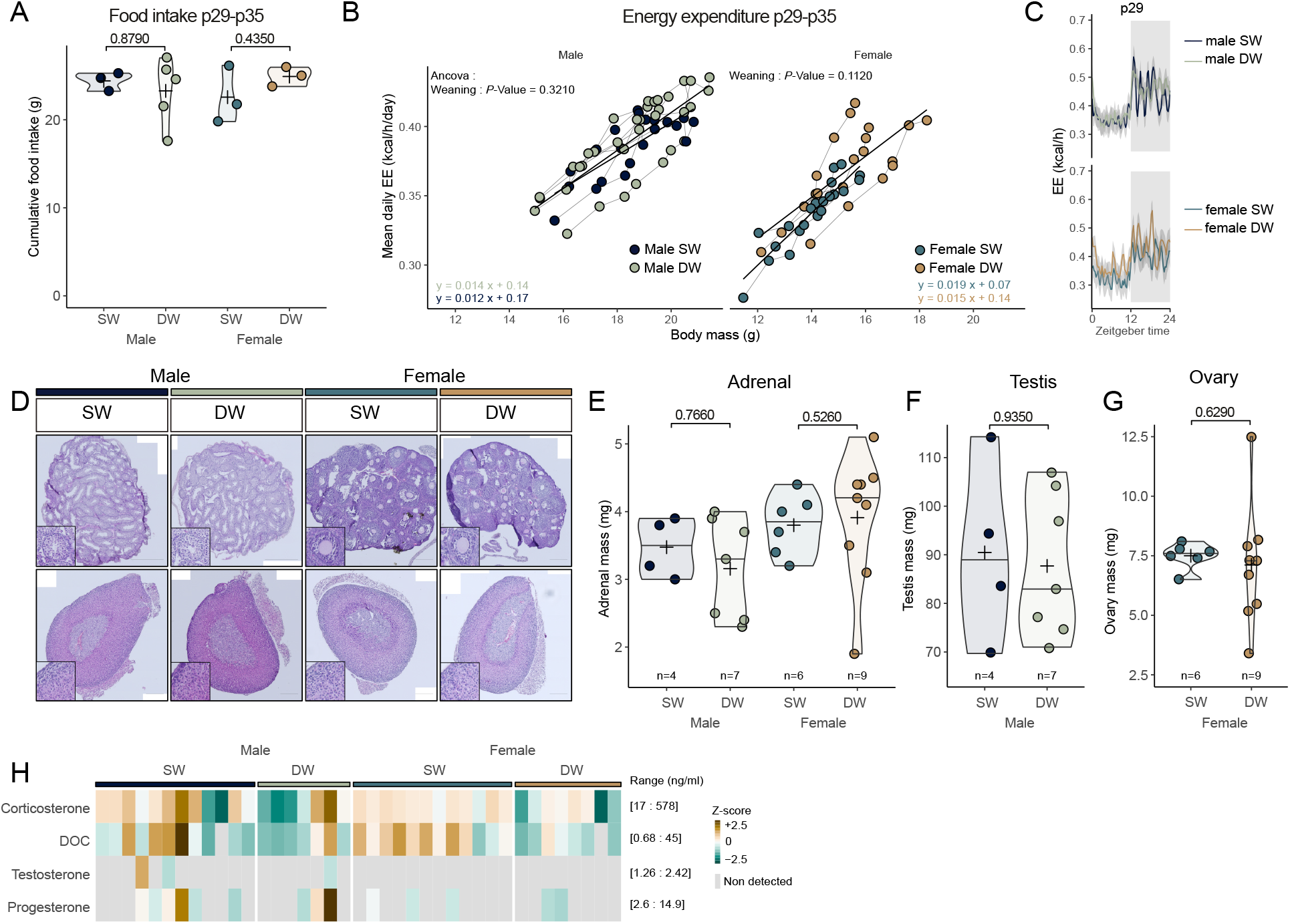
Related to Figure 3. (A) Total food intake between p29 and p35 of SW and DW mice (n = 3-5/group). (B) ANCOVA analysis of energy expenditure as a function of body mass and weaning between p29 and p35 (n = 3-5/group). (C) Energy expenditure at p29 in SW and DW mice. (D) H&E staining of p28 mice’s gonads (top) and adrenal glands (bottom). (E-G) (E) Adrenals. (F) testis and (G) ovary mass of SW and DW mice at p28 (n = 4-9/group). (H) Heatmap representing the plasmatic concentration of DOC, corticosterone, testosterone and progesterone determined by LC–MS/MS in p28 SW and DW mice (n = 7-12/group). *P*-values were determined by two-sided t.test for normally distributed condition or two-sided Mann-Whitney test.

**Figure S3.**
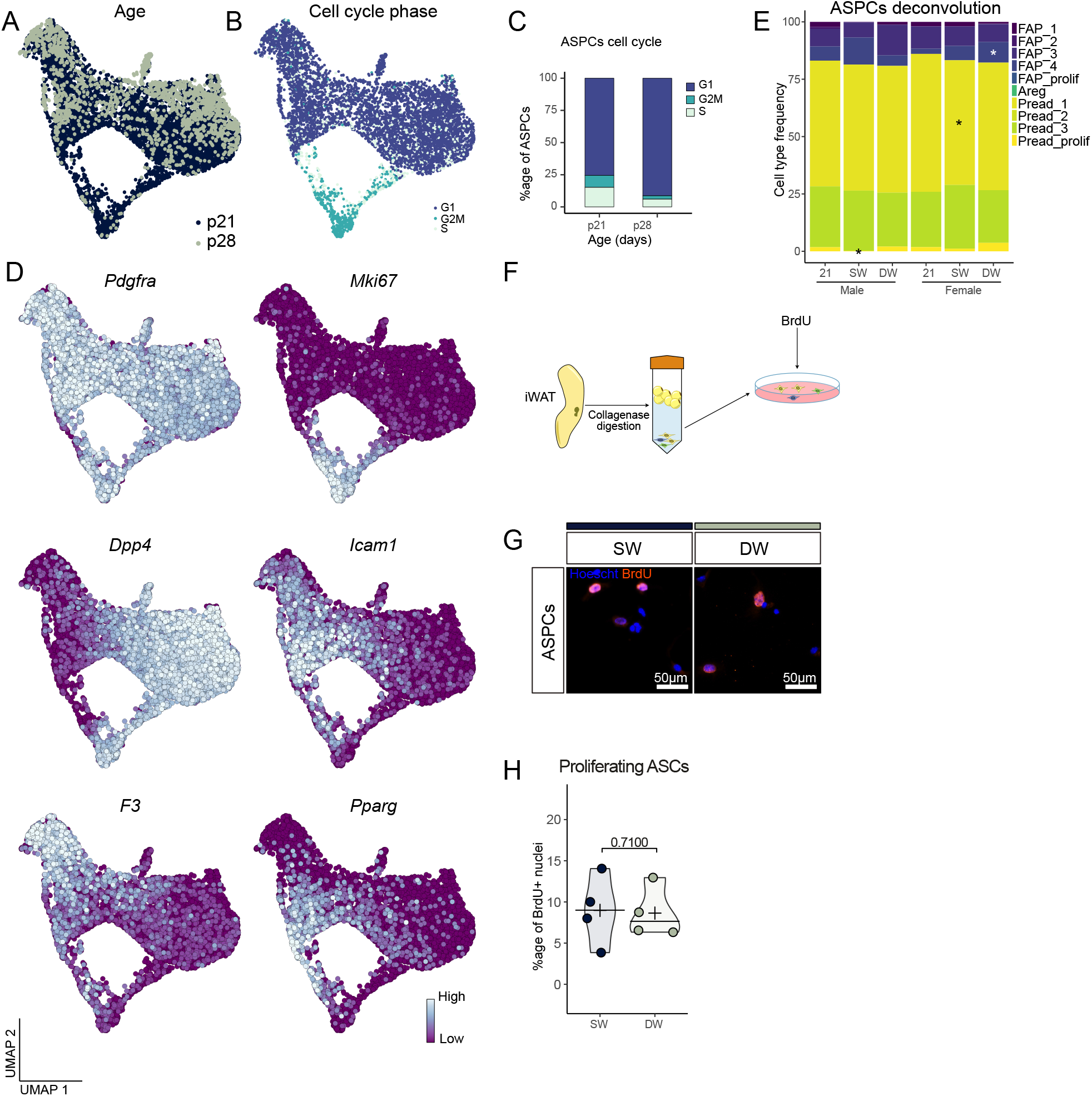
Related to Figure 4. (A) UMAP showing the clustering of p21 and p28 ASPCs (B) UMAP representing ASPCs in G1, G2M or S phase (C) Proportion of ASPCs in G1, G2M or S phase from Qian et al^1^ (D) Feature plots showing expression of selected preadipocytes marker genes. (E) Deconvolution of bulk RNA-seq data with single nuclei cell signature. Asterisks indicate *P*-values of clusters proportion compared to p21, * *P* < 0.05 (F) Simplified scheme of stromal vascular fraction isolation from iWAT. (G and H) BrdU immunofluorescent staining (G) and quantification (H) of proliferating ASPCs isolated from SW or DW male iWAT (n = 4/group). *P*-values were determined by two-sided Mann-Whitney test.

**Figure S4.**
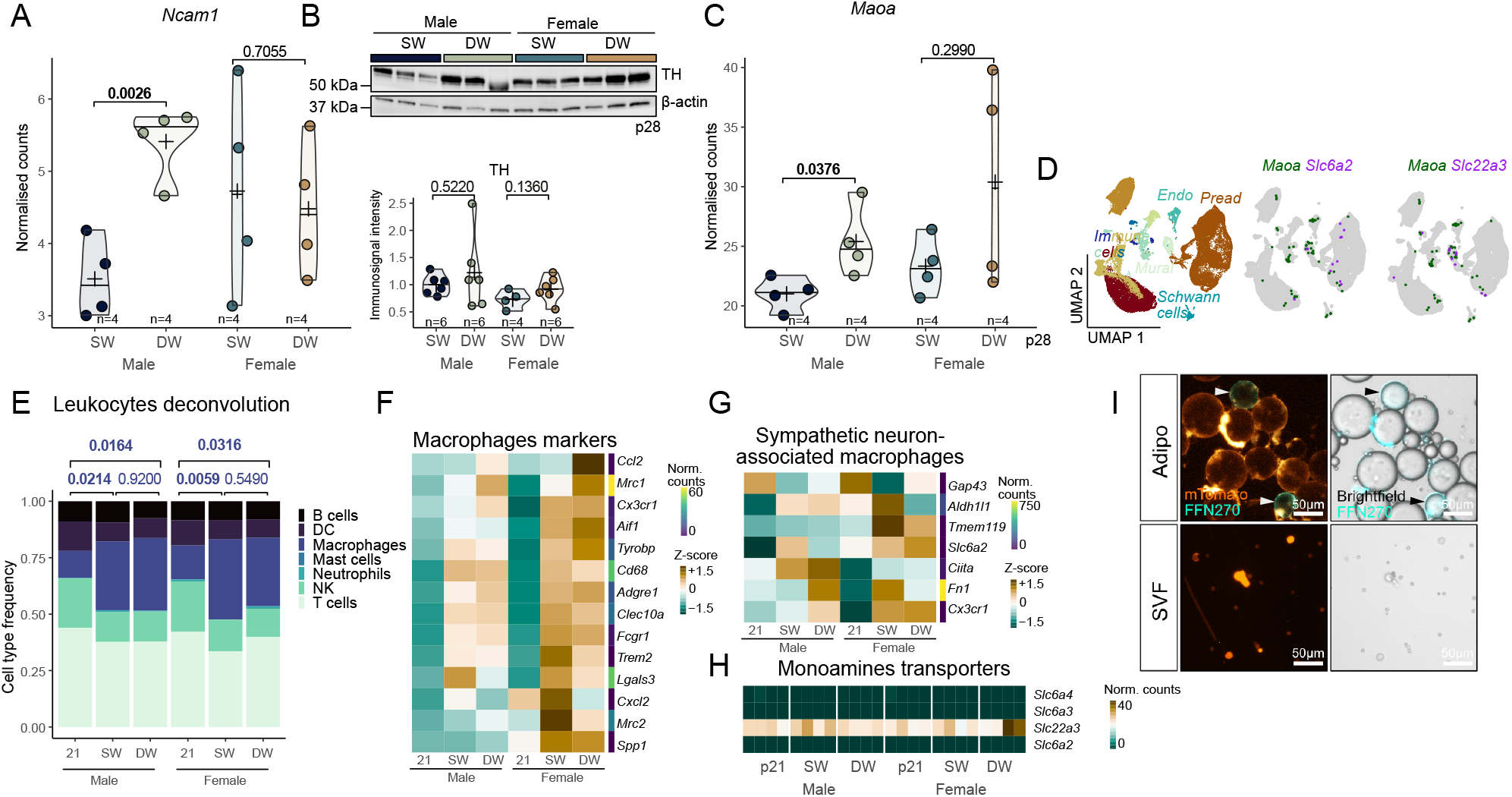
Related to Figure 6. (A) mRNA expression of neuronal marker *Ncam1* in SW and DW iWAT as determined by RNA-seq (n=4). (B) Western blot and quantification of TH in p28 SW and DW iWAT protein extracts (Male SW : n=6, DW : n=6, Female SW : n=4, DW : n=6) (C) mRNA expression of noradrenaline degrading enzyme *Maoa* in SW and DW iWAT as determined by RNA-seq (n=4). (D) UMAP of SVF single cell RNA-seq with coexpression of *Maoa* with *Slc6a2* and *Slc22a3*. (E) Deconvolution of bulk RNA-seq data with single cell immune gene signature. (F) Heatmap showing relative expression of macrophages markers as measured by RNA-seq in p21, p28 SW and p28 DW iWAT (n=4). (G) Heatmap showing relative expression of sympathetic neuron-associated macrophages (SAMs) markers as measured by RNA-seq in p21, p28 SW and p28 DW iWAT (n=4). (H) Heatmap showing absolute expression of monoamines transporters as measured by RNA-seq in p21, p28 SW and p28 DW iWAT (n=4). (I) Rpresentative picture of FFN270 positive cells amongst adipocytes and SVF, membrane are stained by engogenously expressed myristoylated Tomato. Arrows point to positive cells. *P*-values were determined by two-sided Mann-Whitney test.

